# Floral traits predict nectar rewards across diverse plant communities

**DOI:** 10.64898/2026.01.14.699445

**Authors:** E. Herreros-Moya, M. Sinka, H. Portwood, G. MacKay, K. Storer, N. Kühn, K. Willis

## Abstract

Floral nectar is a key energy resource for many animals, yet direct measurement of nectar rewards is time consuming and difficult to apply across taxonomically diverse plant communities. We tested whether certain floral traits can predict nectar rewards across a broad taxonomic and morphological range of flowering plants.

We analysed relationships between floral traits and nectar rewards in 156 species from 42 families. Corolla tube length, corolla base width and corolla shape were recorded alongside nectar volume, sugar concentration and total sugar content, and linear mixed-effects models were used to quantify floral trait–nectar relationships in tubular and non-tubular flowers. Predictive performance was evaluated using observation- and species-level cross-validation, and phylogenetic signal in nectar volume was assessed using Blomberg’s K and Pagel’s λ.

Nectar volume increased strongly with floral dimensions in both tubular and non-tubular flowers, while sugar content was closely associated with nectar volume, with an approximately proportional relationship on the log scale. These relationships support the use of floral traits to predict sugar content, with predictive performance retained for species not included in model fitting (species-level cross-validation R^2^ = 0.43 for tubular and 0.51 for non-tubular flowers). Nectar volume also showed weak but significant phylogenetic signal.

Overall, our findings demonstrate that simple floral traits can provide useful predictors of potential nectar rewards across a taxonomically and morphologically diverse range of flowering plants. Such an approach opens the way for future studies to combine trait-based predictions with information on floral abundance, flowering phenology to investigate how nectar resources vary across landscapes in space and time, with potential applications in pollination ecology and the study of nectar-feeding insects.

## 1. Introduction

Floral nectar plays a fundamental role in plant–animal interactions, providing a critical energy source for many animals and contributing to plant reproductive success through pollination (Kevan & Baker 1983). Most of these interactions occur between plants and insects, where coevolution has shaped a wide diversity of floral traits and interaction strategies, ranging from highly specialised to more generalised systems (Bronstein *et al*. 2006; Duchenne *et al*. 2026; Temeles & Dalsgaard 2026). While some animal pollinators form species-specific associations with plants, many forage opportunistically, selecting floral resources based on availability, quality, and accessibility of the nectar (Irwin *et al*. 2010; van der Kooi *et al*. 2021). Understanding how nectar rewards vary among flowering plants is therefore important for assessing the availability of floral resources and their role in plant– animal interactions.

Nectar rewards are determined by both nectar volume and sugar concentration, which together define the total sugar content available per flower (Corbet 2003). Total sugar content provides a measure of the energetic reward available to nectar-feeding animals, as flowers containing similar nectar volumes may differ in their sugar concentration (Pattrick *et al*. 2025). Although these nectar characteristics can vary considerably among flowers (Farkas & Orosz-Kovács 2003; Galetto & Bernardello 2004), some of this variation may be associated with differences in floral morphology, which can influence how nectar is produced, retained and presented within flowers.

Evidence for these relationships comes from different plant groups and ecological communities. In Mediterranean plant communities, Petanidou and Smets (1995) found that deeper flowers produced more nectar than shallower flowers, while Petanidou *et al*. (2000) identified relationships between nectary structure and nectar secretion in Lamiaceae.

Similar relationships have been reported in other plant groups, with positive associations between corolla length and nectar volume found in *Lonicera implexa* (Lázaro *et al*. 2015), across different species of *Nicotiana* (Kaczorowski et al. 2005), and among hummingbird-visited plants in the Brazilian Atlantic Forest (Castro Tavares *et al*. 2016). These studies suggest potential allometric relationships between floral dimensions and nectar rewards. Shared evolutionary history may also contribute to variation in nectar rewards, as closely related species can share characteristics associated with nectar production. Previous studies have identified phylogenetic signal in nectar traits, suggesting that nectar production may partly reflect shared ancestry (Basnett *et al*. 2025; Keasar & Bodner 2025; Ornelas *et al*. 2007). However, it remains unclear whether floral traits can reliably predict nectar rewards across a broad taxonomic and morphological range of flowering plants.

Establishing such predictive relationships could provide a practical approach when direct nectar measurements are unavailable. Nectar volume is typically quantified by extracting nectar from individual flowers using microcapillary tubes, while sugar concentration is measured using refractometers (Corbet 2003). Although these methods provide direct estimates of nectar rewards, they are time-consuming and logistically challenging when applied across diverse plant communities, particularly when species flower at different times and repeated visits to field sites are impractical. In contrast, many floral dimensions can be measured rapidly and may therefore provide a practical means of estimating potential nectar rewards at the flower level.

In this study, we investigated whether certain floral traits can predict nectar volume and sugar content across a taxonomically and morphologically diverse range of flowering plants. We selected corolla tube length, corolla base width and corolla shape based on previous studies suggesting relationships between these aspects of floral morphology and nectar volume (Castro Tavares *et al*. 2016; Lázaro *et al*. 2015; Romero-Bravo & Castellanos 2024). We hypothesised that nectar volume and sugar content would increase with these floral dimensions and that these relationships could provide a basis for estimating nectar rewards from readily measurable morphological traits.

To test these predictions, we combined floral measurements from 96 species sampled at Oxford Botanic Garden (United Kingdom) and 62 species sampled in natural and semi-natural plant communities in Western Province, Zambia. Together, these datasets comprised 156 species from 42 families, representing a wide range of floral forms and different plant families. We evaluated relationships between floral morphology and nectar rewards using mixed-effects models, examined whether nectar volume could serve as a proxy for total sugar content, tested for differences between sampling locations, and assessed the predictive performance of sugar content models using cross-validation. We also investigated whether nectar volume exhibited phylogenetic signal to explore the extent to which variation in nectar production is associated with shared evolutionary history.

## 2. Materials and Methods

### 2.1. Study sites

The Oxford Botanic Garden (OBG), in Oxford, United Kingdom, spans 1.8 hectares and contains over 5,000 plant species, representing more than 90% of higher plant families. Its phylogenetically arranged order beds include 775 species from 112 flowering plant families. We sampled floral traits from 96 species (31 families) flowering in these beds during July and August 2022. The species were selected to maximise variation in floral morphology and phylogenetic diversity among taxa flowering during the sampling period: selection was therefore constrained by phenology rather than random sampling.

To broaden the taxonomic and morphological range of the dataset, we sampled an additional 62 species from 24 families at field sites in the Kaoma, Nkeyema, and Luampa districts (Western Province, Zambia) in April 2024 (Fig. A1). These sites represent rural landscapes with natural and semi-natural vegetation and form part of a project examining relationships between floral nectar availability and mosquito occurrence (Herreros-Moya et al, 2026 in preparation). In Zambia, species were selected based on an extensive vegetation survey conducted by a local botanist, which identified species collectively accounting for 90% of the total vegetation abundance across the study sites. Floral sampling was restricted to those species that were flowering during the sampling period.

### 2.2. Collection of morphological floral and nectar traits

Floral and nectar traits were measured in the field in both sampling contexts. For each species, 15–20 flowers from different individuals were enclosed in fine mesh bags for 24 hours to prevent insect foraging. Nectar and morphological measurements were subsequently obtained from a subset of these flowers, resulting in 2–15 samples per species. Measurements therefore represent potential nectar production (accumulated secretion) rather than standing crop available to foragers. The following traits were recorded:

#### Corolla shape

Flowers were first categorised as tubular or non-tubular based on the presence or absence of a corolla tube. They were then assigned to morphological categories (campanulate, rotate, funnelform, papilionaceous, bilabiate, tubulate, salverform, or irregular, Fig. A2) following Simpson (2019). Funnelform and campanulate flowers were classified as tubular or non-tubular according to corolla tube development and floral opening angle, with flowers opening wider than 90° considered non-tubular (Wang *et al*. 2020).

#### Corolla size

For tubular flowers, corolla tube length and corolla base width were measured using a 100 mm carbon fibre digital calliper (Fisherbrand 90024-51). For non-tubular flowers, only corolla base width was measured, defined as the width of the floral opening at the base of the corolla.

#### Nectar volume

Nectar volume (µL) was measured using calibrated microcapillary tubes (0.5–5 µL). Volume was calculated as:

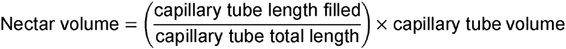

#### Sugar concentration

Sugar concentration was measured as sucrose-equivalent (% w/w) using handheld refractometers (0–50% and 45–80%; Eclipse, Bellingham and Stanley, Tunbridge Wells, UK). A minimum nectar volume of 2 µL was required to obtain a refractometer reading in the field. When individual flowers produced insufficient nectar for this measurement, nectar from multiple flowers of the same species was pooled. For pooled samples, mean values of corolla tube length, corolla base width, and nectar volume were calculated across all flowers included in the pool.

For Asteraceae, measurements were taken at the level of individual florets.

### 2.3. Sugar content per flower

The sugar content per flower represents the total mass of sugar (g) dissolved in the nectar. It was calculated following Corbet (2003) as:

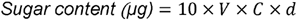

*whereV is* nectar volume (µL),*C* is sugar concentration (% w/w), and *d* is nectar density (mg µL ¹), calculated as:

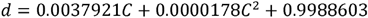

The factor of 10 accounts for conversion of percentage concentration to a proportion and conversion of sugar mass from mg to µg. Sugar content was subsequently converted from µg to g.

### 2.4. Statistical analyses

Relationships between floral traits and nectar rewards were analysed using linear mixed-effects models fitted separately for tubular and non-tubular flowers. Individual flowers, or pooled samples where required for sugar concentration measurements, were treated as the unit of observation, and species identity was included as a random intercept in all models to account for repeated observations within species. This approach allowed us to use all observations while accounting for both within- and between-species variation and unequal numbers of observations among species.

Nectar volume, sugar content, corolla tube length, and corolla base width were log-transformed prior to modelling. Models were compared using likelihood-ratio tests and Akaike Information Criterion (AIC). We first examined relationships between floral traits and nectar volume and sugar content and subsequently evaluated the effect of location (OBG vs Zambia) by comparing the base models with models including location as an additional fixed effect.

All analyses were conducted in R v4.5.1 (R Core Team, 2025).

#### Nectar volume models

For tubular flowers, we modelled nectar volume as a function of corolla tube length, corolla base width, and corolla shape. An interaction between log-transformed corolla tube length and base width was tested but did not improve model fit based on likelihood-ratio tests and AIC and was therefore not retained. The final additive model was:

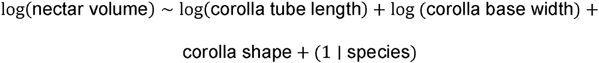

For non-tubular flowers, which lack a defined corolla tube, nectar volume was modelled as a function of corolla base width and corolla shape:

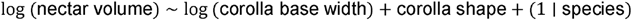

#### Sugar content models

##### Nectar volume as a proxy for sugar content

We evaluated whether nectar volume could serve as a proxy for sugar content when sugar concentration measurements were unavailable. Sugar content per flower is calculated from both nectar volume and sugar concentration (see “Sugar content per flower”); we therefore first assessed the relationships between each component and variation in sugar content.

To further assess the predictive relationship between nectar volume and sugar content, we fitted the following mixed-effects model:

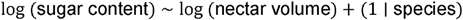

To assess whether floral dimensions explained additional variation in sugar content beyond that accounted for by nectar volume, we compared models including nectar volume and corolla shape with models additionally including corolla tube length and base width.

#### Floral traits as predictors of sugar content

We also modelled sugar content (log-transformed) as a function of floral traits. For tubular flowers, predictors included corolla tube length, corolla base width, and corolla shape:

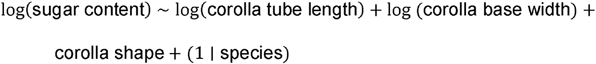

For non-tubular flowers, predictors included corolla base width and corolla shape:

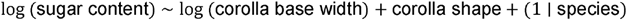

For sugar content analyses, corolla shape categories with very small sample sizes were merged prior to modelling to improve model stability. Campanulate tubular flowers were grouped with funnelform tubular flowers, and irregular non-tubular flowers were grouped with rotate flowers, based on similarities in corolla morphology.

#### Location effects

We examined differences in nectar traits between sampling locations by comparing the base models with models including location (OBG vs Zambia) as an additional fixed effect.

We also tested interactions between location and floral traits to assess whether the relationships between floral traits and nectar rewards differed between locations. As species composition differed between OBG and Zambia, any detected location effects may reflect both environmental differences and species turnover.

#### Model evaluation and sensitivity analyses

Model fit was quantified using marginal and conditional R² values. Multicollinearity among predictors was assessed using variance inflation factors (VIFs).

Predictive performance of the sugar content models was evaluated using Monte Carlo cross-validation (100 iterations; 70/30 training-test splits) (Xu & Liang 2001). We used two cross-validation schemes: observation-level splits, in which individual observations were randomly assigned to training and test sets, and species-level splits, in which entire species were assigned to either the training or test set to evaluate prediction for previously unobserved species. Predictions were based on fixed effects only. Predictive performance was assessed using Pearson correlation (r) between observed and predicted log-transformed sugar content, with R² calculated as the squared correlation.

To assess the robustness of the results to differences in the number of observations among species, we refitted the nectar volume and sugar content models using only species with at least three or four observations.

### 2.5. Phylogenetic signal

Phylogenetic signal in nectar volume was assessed at the species level using mean log-transformed nectar volume for each species to avoid pseudoreplication. Nectar volume was used because measurements were available for substantially more species than sugar content. Phylogenetic signal was quantified using Blomberg’s K (Blomberg *et al*. 2003) and Pagel’s λ (Pagel 1999), implemented in the phytools package in R (Revell 2012).

Both statistics assess whether trait similarity among species is associated with their phylogenetic relatedness. Values approaching zero indicate little phylogenetic signal, whereas values approaching one indicate a pattern more consistent with expectations under a Brownian-motion model of trait evolution.

A dated phylogeny was generated with V.PhyloMaker (Jin & Qian 2019) based on the GBOTB.extended backbone phylogeny (Smith & Brown 2018) and pruned to include only the study taxa. We also repeated the phylogenetic analyses using only species with at least three observations to examine the consistency of the results.

All R code used for nectar volume, sugar content, sensitivity, cross-validation and phylogenetic analyses is archived in Figshare (Appendices S1–S4; see Data Availability Statement).

## 3. Results

A total of 1,077 flowers from 156 species and 42 families were sampled across OBG and Zambia (Table 1), comprising 772 tubular and 305 non-tubular flowers. For each flower, corolla tube length (tubular flowers only), corolla base width, corolla shape, and nectar volume were measured, resulting in over 3,000 individual trait measurements. Floral trait and nectar volume data for all samples are available in Table S1 (see Data Availability Statement).

**Table 1.** Summary of plant samples collected at the OBG and Zambia, including species and families by flower type.

|  | **TOTAL** |  |  | **Tubular** |  |  | **Non-Tubular** |  |  |
|---|---|---|---|---|---|---|---|---|---|
|  | **Samples** | **Species** | **Families** | **Samples** | **Species** | **Families** | **Samples** | **Species** | **Families** |
| **OBG** | 815 | 96 | 31 | 603 | 71 | 21 | 212 | 25 | 11 |
| **Zambia** | 262 | 62 | 24 | 169 | 40 | 17 | 93 | 23 | 11 |
| **TOTAL** | 1077 | 156* | 42 | 772 | 108* | 38 | 305 | 48 | 22 |

\* The total does not equal OBG + Zambia because some species occurred in both locations.

Sugar concentration was measured for a subset of 183 samples from 102 species, as a minimum nectar volume of 2 µL was required to obtain a refractometer reading (Table A1). The full sugar content dataset, including all species for which sugar concentration was measured, is available in Table S2 (see Data Availability Statement).

### 3.1. Nectar volume models

#### Tubular flowers

Analysis of 772 tubular flowers from 108 species showed that both corolla tube length and base width were strong positive predictors of nectar volume (both p < 0.001; Fig. 1A), with the model explaining a substantial proportion of the variation (marginal R² ≈ 0.47; conditional R² ≈ 0.81).

**Fig. 1.**
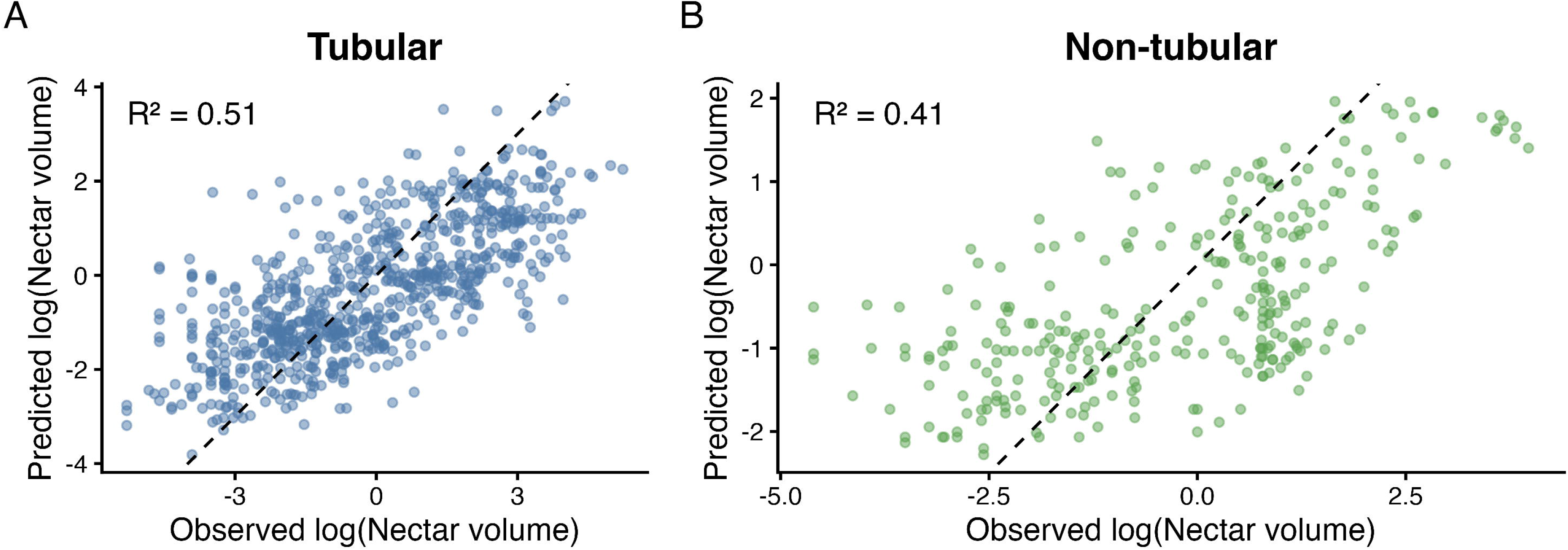
Observed versus predicted nectar volume based on mixed-effects models. **(A)** Tubular flowers. **(B)** Non-tubular flowers. Points represent individual flowers, and the dashed line indicates the 1:1 relationship between observed and predicted values. Predictions are based on fixed effects only.

Corolla shape also contributed to variation in nectar volume (*p* = 0.019), although differences among shapes were small relative to the associations with corolla tube length and base width (Fig. A3).

Including location (OBG vs Zambia) improved model fit (χ² = 4.48, df = 1, p = 0.034), with lower nectar volumes in Zambia. However, interactions between location and floral dimensions did not improve model fit (χ² = 3.60, df = 2, p = 0.166).

#### Non-tubular flowers

Analysis of 305 non-tubular flowers from 48 species showed that corolla base width was a strong positive predictor of nectar volume (p < 0.001; Fig. 1B), with the model explaining a moderate proportion of the variation (marginal R² ≈ 0.34; conditional R² ≈ 0.74).

Corolla shape did not contribute significantly to variation in nectar volume after accounting for corolla width (*p* = 0.253; Fig. A3).

Including location improved model fit (χ² = 9.89, df = 1, p = 0.0017), but the interaction between location and corolla width did not (χ² = 0.16, df = 1, p = 0.69).

### 3.2. Sugar content models

#### Nectar volume as a proxy for sugar content

Sugar content showed a strong positive relationship with nectar volume (Fig. 2A). On the log scale, the relationship was approximately proportional (β ≈ 1), indicating proportional scaling between nectar volume and sugar content. In contrast, sugar concentration showed a much weaker relationship with total sugar content (Fig. 2B). Adding floral dimensions to models containing nectar volume and corolla shape further improved model fit (χ² = 8.99, df = 2, p = 0.011).

**Fig. 2.**
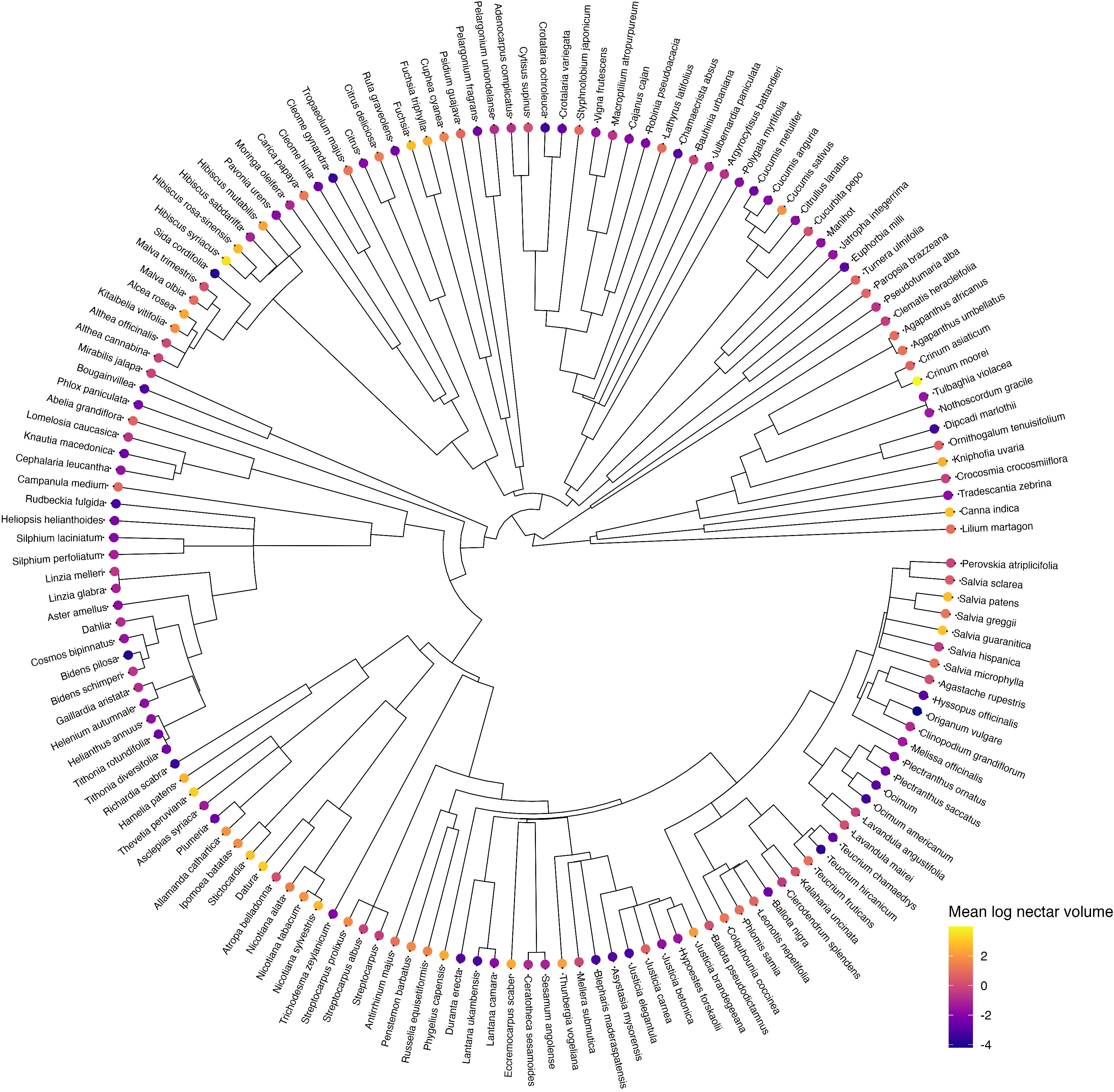
Relationships between sugar content per flower and **(A)** nectar volume (µL) and **(B)** sugar concentration (%). Points represent individual samples and lines represent fitted relationships.

#### Floral traits as predictors of sugar content

We next evaluated whether floral traits could predict sugar content in the absence of nectar volume measurements.

For tubular flowers (129 samples from 72 species), both corolla tube length and base width were strong positive predictors of sugar content (both p < 0.001; Fig. 3A), with the model explaining a substantial proportion of the variation (marginal R² ≈ 0.50; conditional R² ≈ 0.77). Corolla shape contributed relatively little to variation in sugar content (Fig. 4A). Including location did not improve model fit (χ² = 2.43, df = 1, p = 0.12), and interactions between location and floral traits were not significant (p = 0.85).

**Fig. 3.**
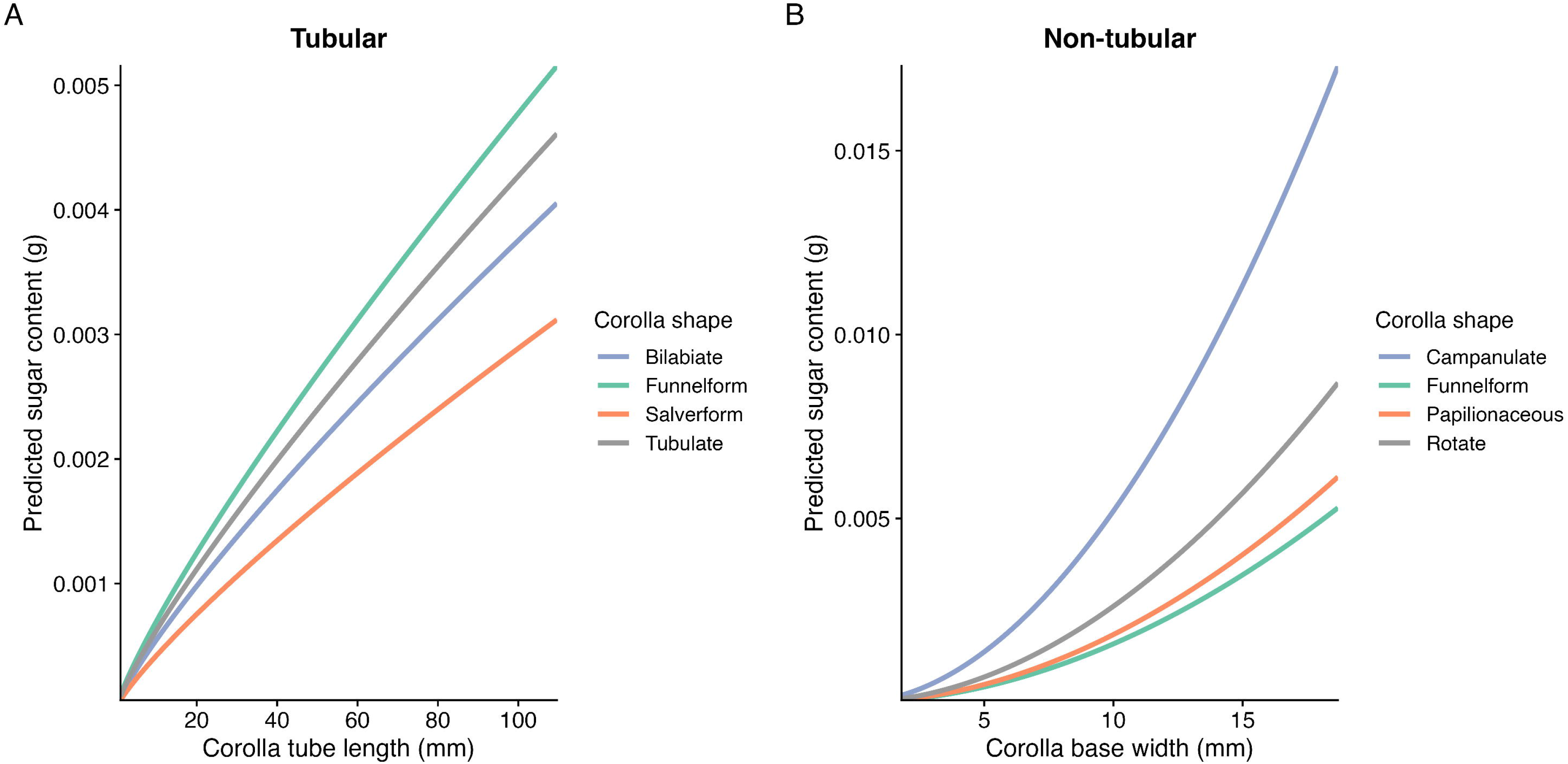
Observed versus predicted sugar content per flower based on mixed-effects models. **(A)** Tubular flowers. **(B)** Non-tubular flowers. Points represent individual samples, and the dashed line indicates the 1:1 relationship between observed and predicted values. Predictions are based on fixed effects only.

**Fig. 4.**
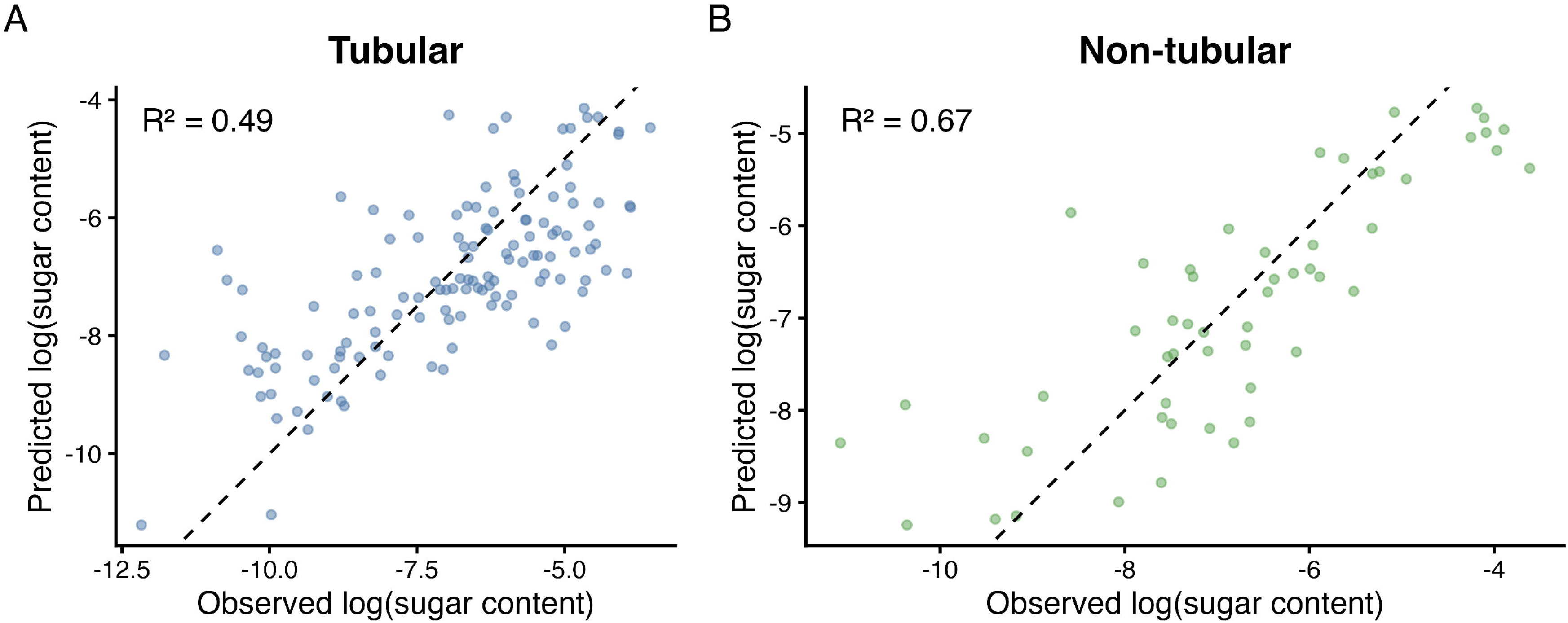
Predicted sugar content per flower as a function of floral traits. **(A)** Tubular flowers: sugar content as a function of corolla tube length. **(B)** Non-tubular flowers: sugar content as a function of corolla base width. Lines represent predictions from mixed-effects models for different corolla shapes, with other model predictors held constant.

For non-tubular flowers (54 samples from 30 species), corolla base width was a strong positive predictor of sugar content (β ≈ 1.9, p < 0.001; Fig. 3B), with the model explaining a substantial proportion of the variation (marginal R² ≈ 0.61; conditional R² ≈ 0.80). Corolla shape contributed relatively little compared with corolla base width (Fig. 4B). Including location improved model fit (χ² = 5.49, df = 1, p = 0.019), but interactions between location and floral traits were not significant (p = 0.65).

Fixed-effect estimates for all mixed-effects models are provided in the Appendix (Tables A2–A5).

### 3.3. Model performance and robustness

Monte Carlo cross-validation indicated moderate predictive performance for the sugar content models. For tubular flowers, mean R² was 0.48 for observation-level splits and 0.43 for species-level splits, in which predictions were made for species not included in model fitting. For non-tubular flowers, mean R² was 0.63 and 0.51, respectively.

Sensitivity analyses restricting the datasets to species with at least three and four observations produced broadly consistent patterns in the principal floral-trait effects (Tables A6 and A7), although estimates became less precise as sample sizes decreased.

### 3.4. Phylogenetic signal

Phylogenetic analyses were conducted using species mean log-transformed nectar volume across 154 taxa from 42 families (Fig. 5). Three *Fuchsia* species (*Fuchsia rivendell*, *Fuchsia checkerboard* and *Fuchsia thamar*) were grouped at the genus level because species-level phylogenetic matches were unavailable.

**Fig. 5.**
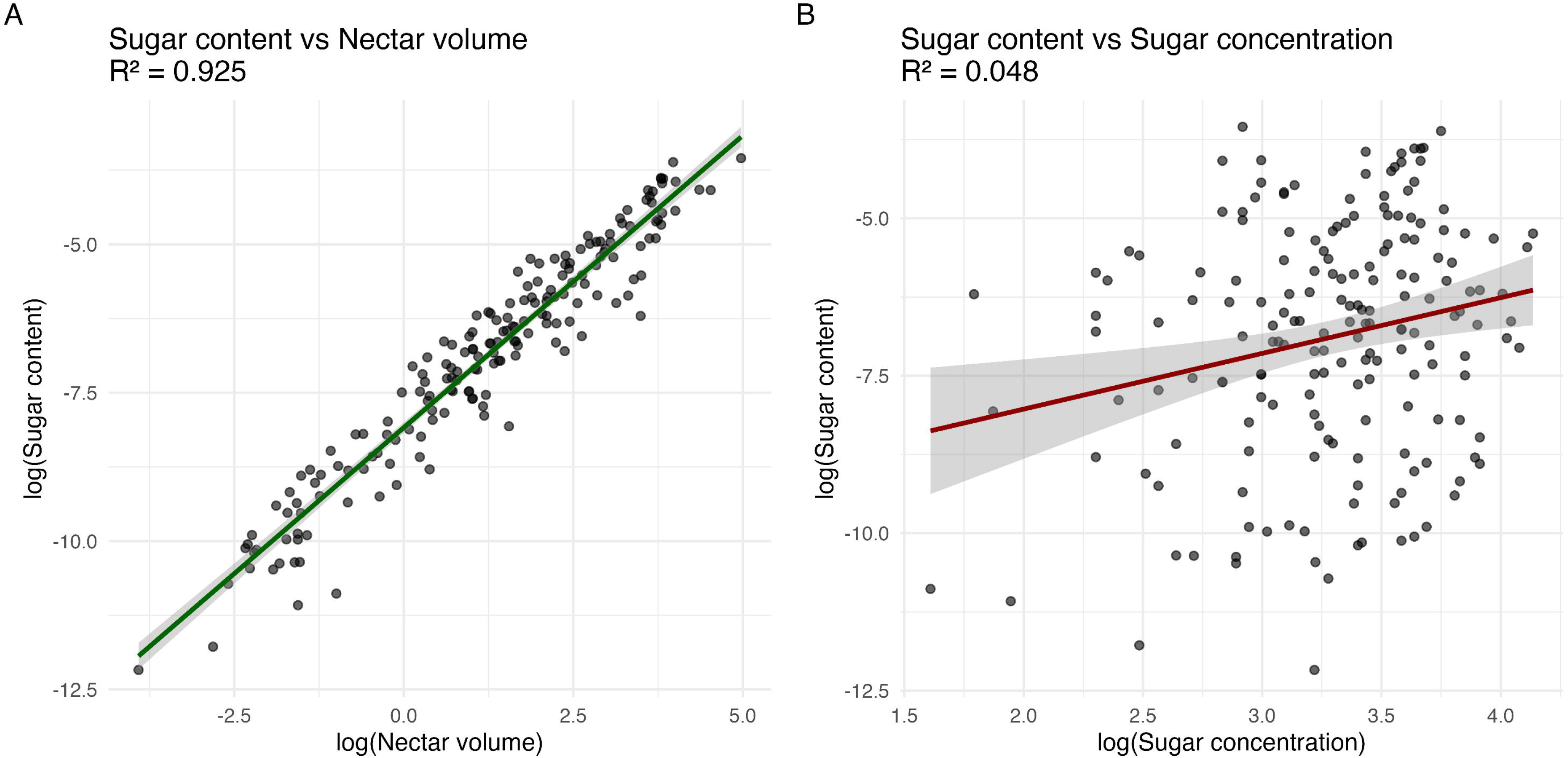
Dated phylogeny of 154 flowering plant taxa showing the distribution of mean log-transformed nectar volume across lineages. Tip colours represent species-level mean log-transformed nectar volume. Nectar volume showed weak but significant phylogenetic signal (Blomberg’s K = 0.19, *P* = 0.001; Pagel’s λ = 0.31, *P* = 0.006).

Nectar volume showed weak but significant phylogenetic signal (Blomberg’s K = 0.19, P = 0.001; Pagel’s λ = 0.31, P = 0.006), indicating relatively weak phylogenetic structuring of nectar volume across the sampled species.

Results were consistent when restricting the analysis to species with at least three observations (K = 0.22, P = 0.001; λ = 0.47, P < 0.001).

## 4. Discussion

Our results show that floral traits provide useful predictors of nectar rewards across taxonomically and morphologically diverse flowering plants. Across both tubular and non-tubular flowers, measures of corolla size explained substantial variation in nectar volume. Sugar content per flower was strongly associated with nectar volume, suggesting that differences in sugar reward were related more to the amount of nectar produced than to variation in sugar concentration. Although species identity accounted for additional variation, floral dimensions remained strong predictors of nectar rewards.

Nectar production was associated with floral size across both floral types. In tubular flowers, nectar volume increased with both corolla tube length and base width, with larger and longer tubes containing more nectar. These results are consistent with previous studies showing that nectar production scales with floral size (Lázaro *et al*. 2015; Ornelas *et al*. 2007; Petanidou & Smets 1995). Corolla shape also contributed to variation in nectar volume, but differences among shapes were small compared with the effects of corolla tube length and width. A similar pattern was reported by Keasar and Bodner (2025), who found that corolla tube length had a greater influence on nectar sugar production than floral symmetry.

In non-tubular flowers, nectar volume increased with corolla base width, while corolla shape contributed relatively little after accounting for width. Floral size therefore captures a substantial component of the variation in nectar volume across both floral types, while the contribution of corolla shape appears more limited.

Importantly, the sugar content models retained moderate predictive performance when applied to species not included in model fitting, suggesting that relationships between floral traits and nectar rewards extend beyond the species used to develop the models. This is particularly relevant for field surveys involving species for which nectar measurements are unavailable, as floral dimensions can be measured relatively quickly and with simple equipment.

Nectar rewards also varied between the two sampling locations. Nectar volume differed between OBG and Zambia for both tubular and non-tubular flowers, while sugar content differed between locations for non-tubular flowers. We believe that differences in environmental conditions, sampling periods and species composition may have contributed to this variation. However, we did not find significant interactions between location and floral traits for either nectar volume or sugar content, suggesting that the relationships between floral traits and nectar rewards did not differ significantly between OBG and Zambia. Further studies across different locations and sampling periods would help to determine whether these relationships remain consistent across environments and over time.

We also found a weak but significant phylogenetic signal in nectar volume, indicating that closely related species resemble one another more than expected by chance, but that this trait is only weakly conserved across the sampled species. This suggests that shared evolutionary history contributes to variation in nectar production, although other factors, including ecological conditions and floral characteristics, also play important roles. Similar patterns of weak or absent phylogenetic signal in nectar traits have been reported in previous studies (Basnett *et al*. 2025; Lanuza *et al*. 2023; Ortiz *et al*. 2021), while other studies have identified phylogenetic influences on nectar production (Keasar & Bodner 2025; Ornelas *et al*. 2007). These differences may reflect variation among studies in taxonomic scale, floral diversity and ecological context, as nectar production may be influenced by responses to pollination environments as well as shared ancestry.

Predicting how much nectar a flower contains, however, is only one part of understanding its value as a resource. Floral morphology may also influence which insects are able to access that nectar. For example, in tubular flowers, longer and narrower corollas may prevent insects with short proboscides from reaching the nectar, while wider or more open flowers may be accessible to a greater variety of insects (Lázaro *et al*. 2015). Such physical limitations can act as mechanical filters that shape which resources different insect species can access (Castellanos *et al*. 2004; Harder 1983; Martins & Johnson 2013; Torres & Galetto 2002). Our approach therefore predicts potential nectar rewards rather than the amount available to any particular insect, as accessibility also depends on mouthpart morphology, particularly proboscis length and shape, and feeding behaviour. Some nectar-feeding insects can also bypass these barriers by piercing floral tissues, making the relationship between floral morphology and nectar accessibility more complex. Combining predictions of nectar rewards with information on floral accessibility could help us better understand which resources are available to different insect groups.

This distinction may be particularly relevant for nectar-feeding insects that are less frequently considered in studies of floral resources, including mosquitoes. Plant sugars are an important energy source for mosquitoes, influencing their survival, activity and reproductive performance (Foster 1995; Gary Jr & Foster 2004; Gouagna *et al*. 2014; Impoinvil *et al*. 2004; Manda *et al*. 2007; Müller *et al*. 2017; Müller *et al*. 2011). The distribution and accessibility of floral sugar resources may therefore have important implications for mosquito ecology. In this context, trait-based predictions of floral sugar rewards could help investigate how variation in plant sugar availability relates to mosquito populations and their use of plant communities. These predictions could be used to investigate whether variation in floral sugar availability is linked to vector-borne disease dynamics (Gu *et al*. 2011; Stone *et al*. 2012). Establishing these links, however, would require combining estimates of floral resources with information on mosquito feeding behaviour, accessibility and population dynamics.

At a broader scale, our findings also have practical applications for quantifying nectar resources across diverse plant communities. Previous studies in the UK have shown how estimates of nectar resources can be used to investigate variation in floral sugar availability across space and time (Baude *et al*. 2016; Hicks *et al*. 2016; Tew *et al*. 2021). Building on this work, trait-based predictions could facilitate assessments of floral sugar resources across landscapes by reducing the need for direct nectar measurements. Flower-level predictions could then be combined with information on floral abundance and phenology to estimate how nectar sugar resources vary across communities and through time. Such approaches could be extremely useful in pollination ecology, conservation and studies of animal populations that depend on floral resources, especially when studying how nectar availability may change with shifts in flowering phenology caused by climate change (Takkis *et al*. 2018) and changes in plant-community composition, including those resulting from invasive species (Agha *et al*. 2021; Müller *et al*. 2017; Nyasembe *et al*. 2015; Sinka *et al*. 2020).

There are some limitations to consider when interpreting our results. First, nectar measurements represent accumulated secretion over 24 hours rather than the standing crop available to foragers at any particular time. Second, nectar production can vary within species in response to environmental conditions, flower age and other temporal factors that we did not incorporate into our models. Measurements also represent particular sampling periods at OBG and in Zambia and may not capture temporal variation within either location. Third, sugar concentration could only be measured in samples with sufficient nectar for refractometer readings, which meant that fewer observations were available for the sugar content analyses. Finally, the number of observations differed among species, but when we repeated the analyses using only species with at least three or four observations, we found similar relationships between floral traits and nectar rewards.

In summary, corolla dimensions capture meaningful variation in nectar rewards across a wide range of flowering plants. Corolla size was consistently associated with nectar volume, which in turn was closely associated with total sugar reward. This provides a practical approach for estimating potential nectar rewards from floral traits and opens the possibility of combining these predictions with data on floral abundance, phenology, and accessibility to better understand how nectar resources vary across plant communities and landscapes.

## 5. Acknowledgements

We thank Mundia Masuzyo and Frank Ndalama at PATH Kaoma, Zambia, for their invaluable support in liaising with local communities during flower sampling, as well as the entire PATH Kaoma team and all volunteers who allowed access to their gardens for fieldwork. We also thank the Oxford Botanic Garden for supporting field measurements during the summer months.

This study was funded by the Innovative Vector Control Consortium (IVCC) with support from the Bill & Melinda Gates Foundation (grant INV-007509), the Swiss Agency for Development and Cooperation (SDC; grant 81067480), and UK Aid (grant 30041-105). The findings and conclusions presented here are those of the authors and do not necessarily reflect the positions or policies of the funding organisations.

## 6. Author contributions

EHM conceived and designed the study. EHM, HP, JM and KS collected the data. NK contributed to model development. EHM performed the statistical analyses and modelling and wrote the first draft of the manuscript. All authors contributed to manuscript revision and approved the final version. KW secured funding, and KW and MS provided overall supervision.

## 8. Data availability statement

Nectar volume and sugar content datasets and all R code required to reproduce the analyses and figures are archived in Figshare: https://doi.org/10.6084/m9.figshare.32346678.

## Appendix

**Table A1.**
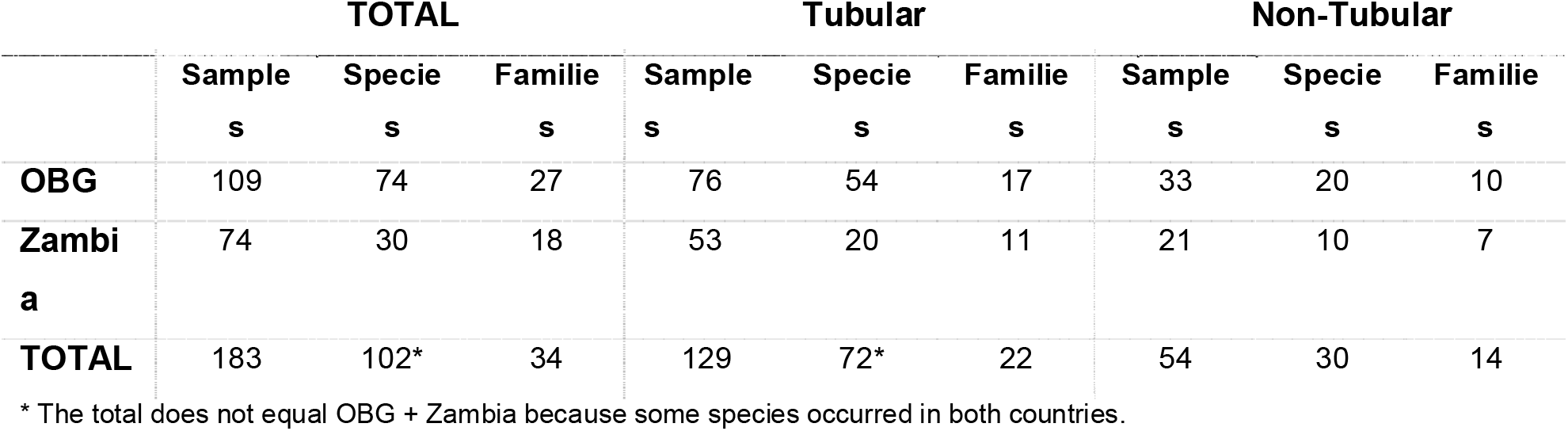
Summary of samples with sugar concentration measurements by flower type, species, and family at each location.

|  | **TOTAL** |  |  | **Tubular** |  |  | **Non-Tubular** |  |  |
|---|---|---|---|---|---|---|---|---|---|
|  | **Samples** | **Species** | **Families** | **Samples** | **Species** | **Families** | **Samples** | **Species** | **Families** |
| **OBG** | 109 | 74 | 27 | 76 | 54 | 17 | 33 | 20 | 10 |
| **Zambia** | 74 | 30 | 18 | 53 | 20 | 11 | 21 | 10 | 7 |
| **TOTAL** | 183 | 102* | 34 | 129 | 72* | 22 | 54 | 30 | 14 |

\* The total does not equal OBG + Zambia because some species occurred in both countries.

**Table A2.** Fixed-effect estimates from the linear mixed-effects model predicting nectar volume in tubular flowers.

| **Predictor** | **Estimate** | **SE** | **t-value** | **95% CI** |
|---|---|---|---|---|
| log_length | 1.34 | 0.12 | 10.93 | (1.10, 1.58) |
| log_width | 0.52 | 0.13 | 4.02 | (0.27, 0.77) |
| Campanulate | 0.01 | 0.57 | 0.02 | (−1.13, 1.15) |
| Funnelform | 0.62 | 0.38 | 1.64 | (−0.13, 1.38) |
| Salverform | −0.74 | 0.42 | −1.78 | (−1.57, 0.09) |
| Tubulate | 0.61 | 0.32 | 1.90 | (−0.03, 1.24) |
| Marginal R<sup>2</sup> = 0.47; Conditional R<sup>2</sup> = 0.81 |  |  |  |  |

**Table A3.** Fixed-effect estimates from the linear mixed-effects model predicting nectar volume in non-tubular flowers.

| Predictor | Estimate | SE | t-value | 95% CI |
|---|---|---|---|---|
| log_width | 1.16 | 0.30 | 3.87 | (0.57, 1.75) |
| Funnelform | −0.29 | 0.73 | −0.39 | (−1.77, 1.20) |
| Irregular | −1.55 | 0.79 | −1.97 | (−3.14, 0.03) |
| Papilionaceous | −1.29 | 0.65 | −1.97 | (−2.60, 0.02) |
| Rotate | −1.05 | 0.62 | −1.71 | (−2.29, 0.18) |
| Marginal R<sup>2</sup> = 0.34; Conditional R<sup>2</sup> = 0.74 |  |  |  |  |

**Table A4.**
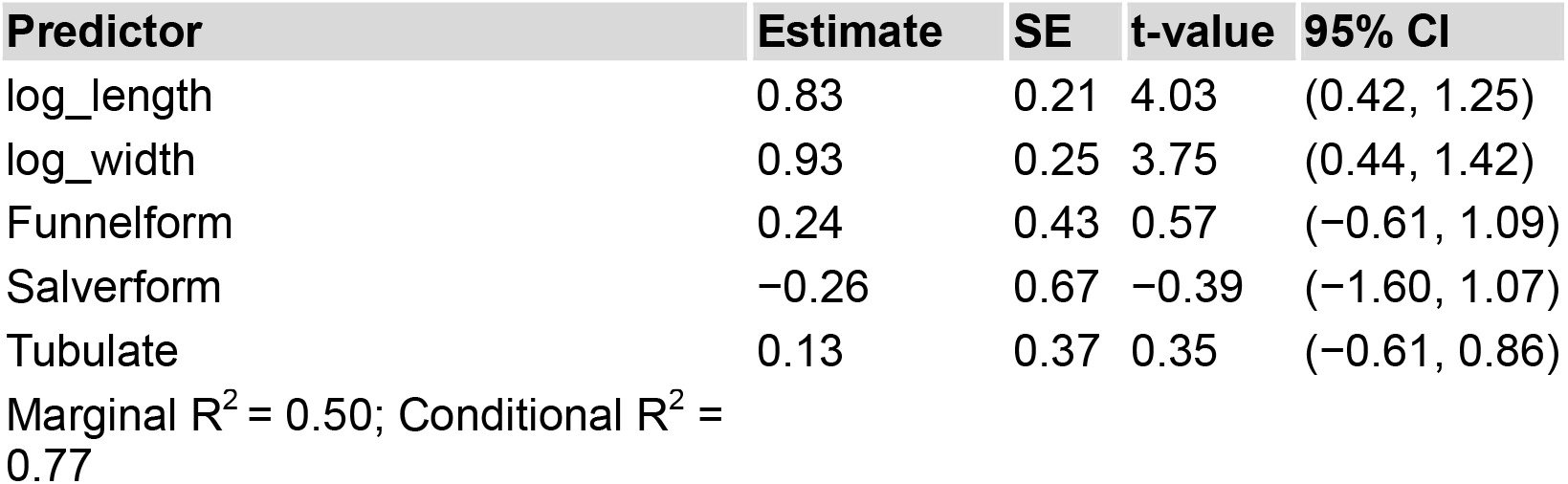
Fixed-effect estimates from the linear mixed-effects model predicting sugar content in tubular flowers.

**Table A5.**
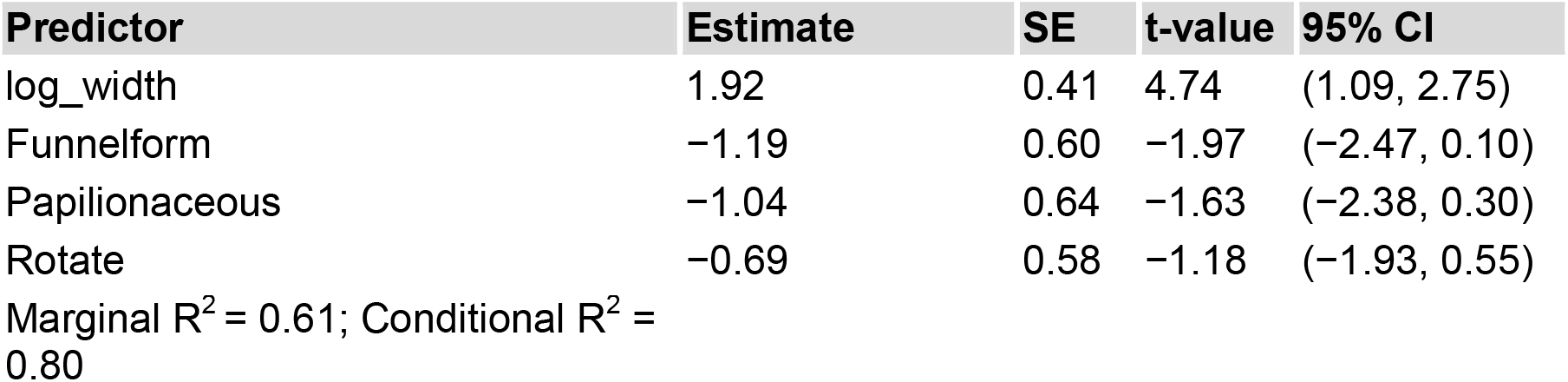
Fixed-effect estimates from the linear mixed-effects model predicting sugar.

**Table A6.** Sensitivity analyses of mixed-effects models assessing the effects of floral traits on nectar volume. Models were refitted using subsets of species with at least three and four.

| **Tubular flowers** |  |  |  |
|---|---|---|---|
| **Predictor** | **Full (n≥1)** | **≥3 obs** | **≥4 obs** |
| log_length | 1.34 | 1.31 | 1.29 |
| log_width | 0.52 | 0.53 | 0.54 |
| **Non-tubular flowers** |  |  |  |
| **Predictor** | **Full (n≥1)** | **≥3 obs** | **≥4 obs** |
| log_width | 1.16 | 1.12 | 1.11 |
| **Model fit** | **Full (n≥1)** | **≥3 obs** | **≥4 obs** |
| Tubular R²m<sup>*</sup> | 0.47 | 0.47 | 0.46 |
| Tubular R²c<sup>*</sup> | 0.81 | 0.82 | 0.82 |
| Non-tubular R²m<sup>*</sup> | 0.34 | 0.40 | 0.41 |
| Non-tubular R²c<sup>*</sup> | 0.74 | 0.73 | 0.73 |

<sup>\*</sup>R²m, marginal R²; R²c, conditional R².

**Table A7.** Sensitivity analyses of mixed-effects models assessing the effects of floral traits on sugar content. Models were refitted using subsets of species with at least three and four.

| **Tubular flowers** |  |  |  |
| **Predictor** | **Full (n≥1)** | **≥3 obs** | **≥4 obs** |
|---|---|---|---|
| log_length | 0.83 | 1.14 | 0.99 |
| log_width | 0.93 | 0.18 | −0.68 |
| **Non-tubular flowers** |  |  |  |
| **Predictor** | **Full (n≥1)** | **≥3 obs** | **≥4 obs** |
| log_width | 1.92 | 2.08 | 2.15 |
| **Model fit** | **Full (n≥1)** | **≥3 obs** | **≥4 obs** |
| Tubular R²m<sup>*</sup> | 0.50 | 0.27 | 0.40 |
| Tubular R²c<sup>*</sup> | 0.77 | 0.72 | 0.62 |
| Non-tubular R²m<sup>*</sup> | 0.61 | 0.73 | 0.78 |
| Non-tubular R²c<sup>*</sup> | 0.80 | 0.73 | 0.78 |

<sup>\*</sup>R²m, marginal R²; R²c, conditional R².

**Fig. A1.**
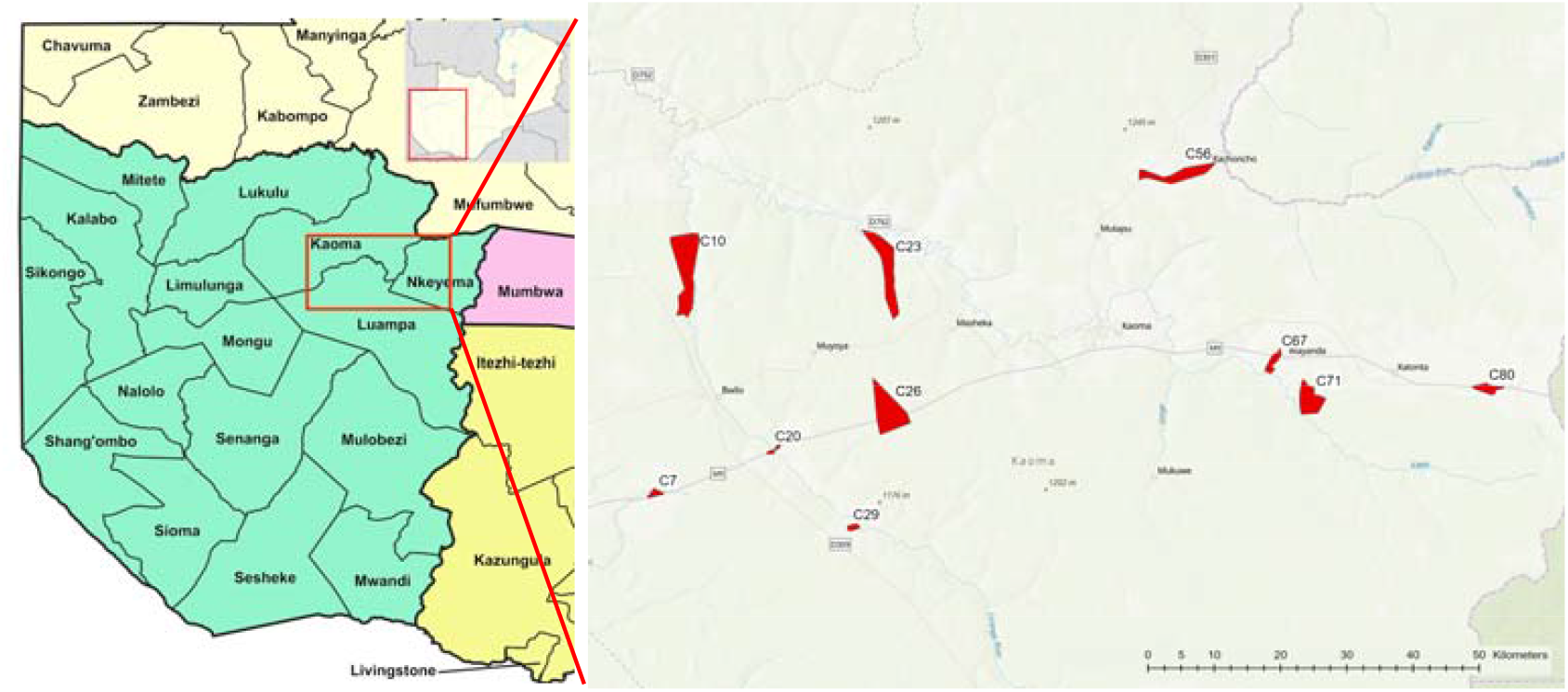
Map of study sites in Kaoma, Nkeyema and Luampa districts, Western Province, Zambia. Red polygons indicate the locations and approximate areas of the field clusters where floral trait and nectar data were collected. Base map adapted from “Zambia districts in Western Province 2022” (Wikimedia Commons).

**Fig. A2.**
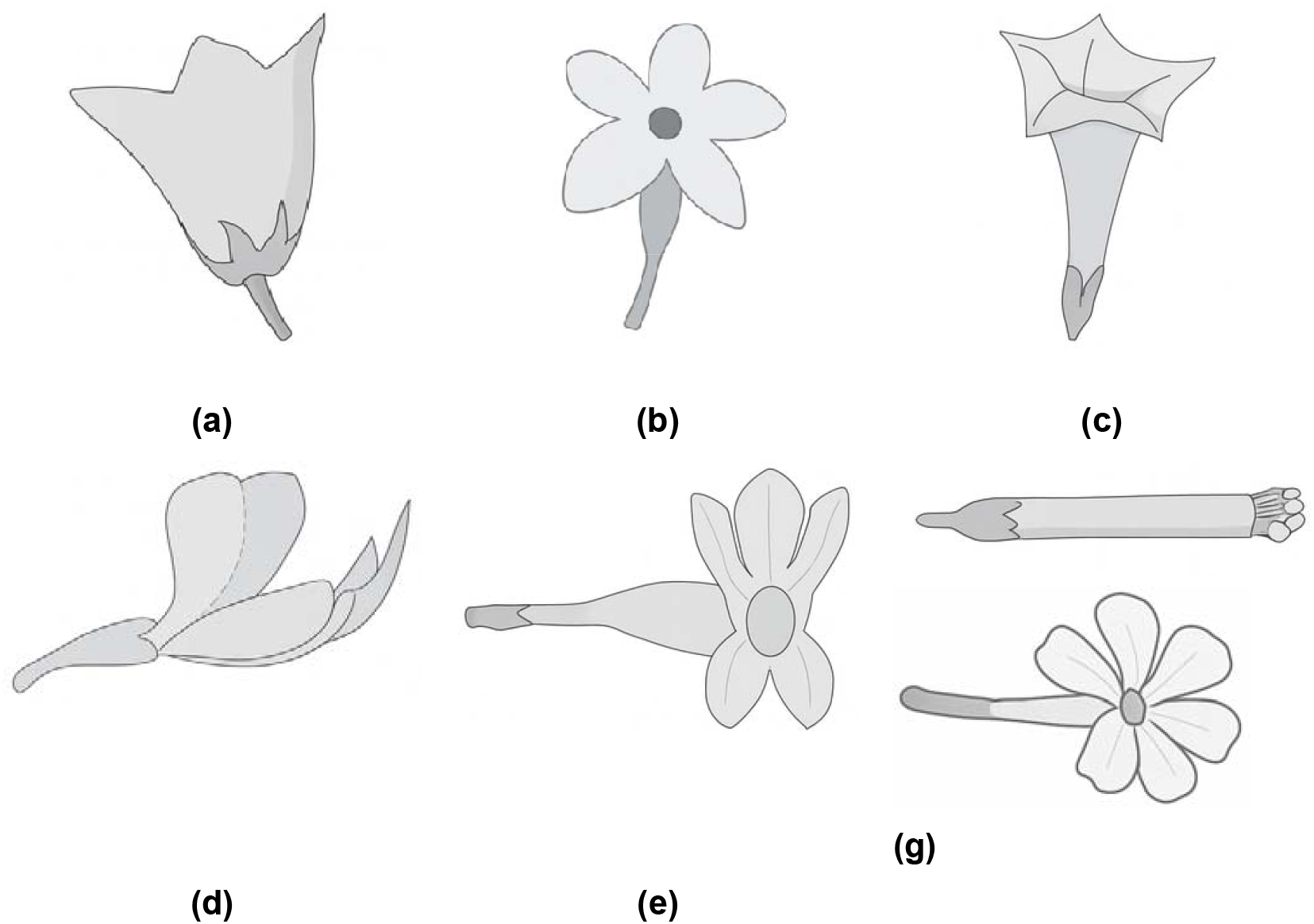
Corolla shapes illustrating floral morphological diversity: **(a)** campanulate, **(b)** rotate, **(c)** funnelform, **(d)** papilionaceous, **(e)** bilabiate, **(f)** tubulate, and **(g)** salverform.

**Fig. A3.**
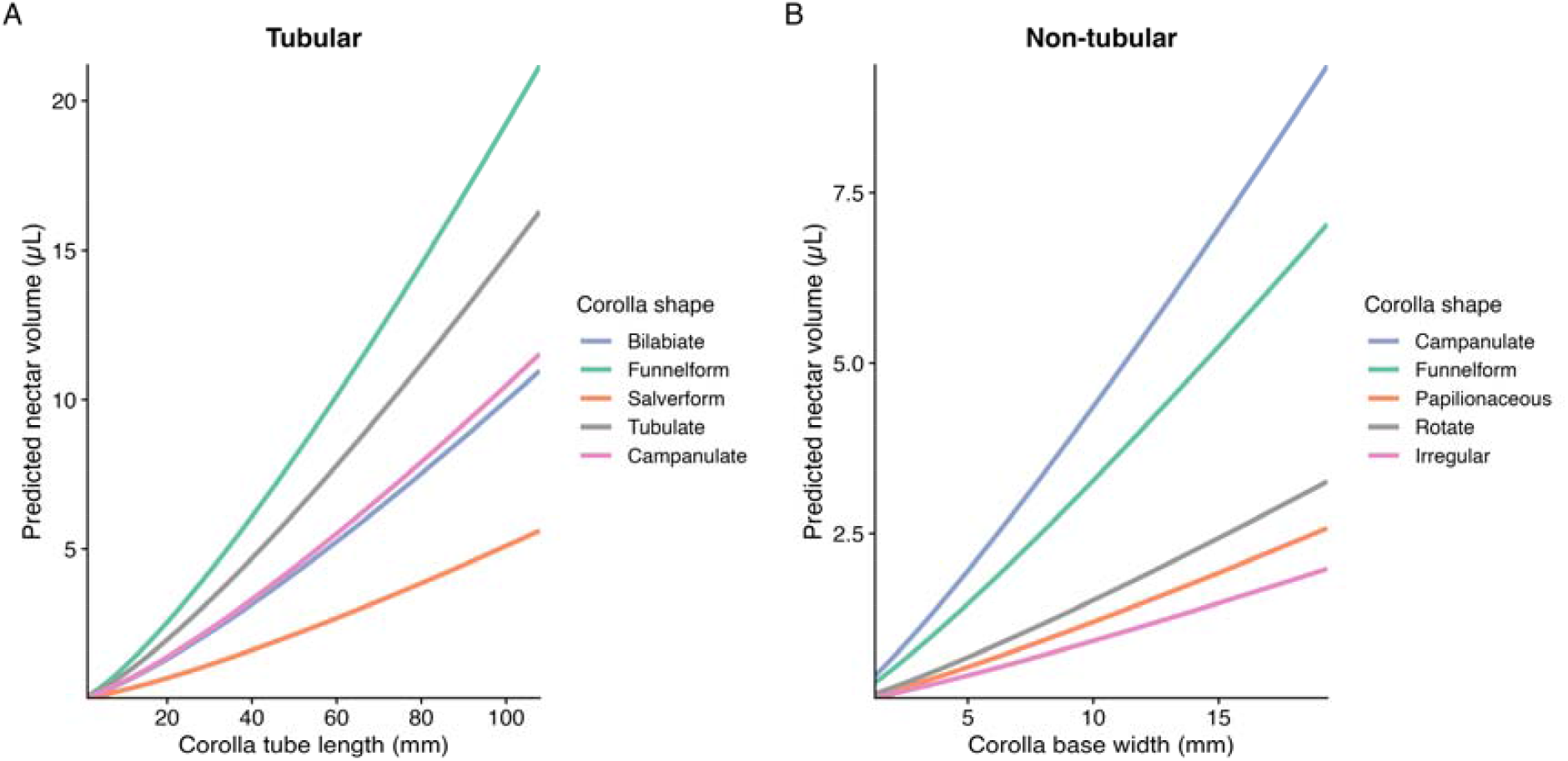
Predicted nectar volume as a function of floral traits. **(A)** Tubular flowers: nectar volume as a function of corolla tube length. **(B)** Non-tubular flowers: nectar volume as a function of corolla base width. Lines represent mixed-effects model predictions for different corolla shapes, with other model predictors held constant.

## References

Agha, S.B., Alvarez, M., Becker, M., Fèvre, E.M., Junglen, S. & Borgemeister, C. (2021). Invasive Alien Plants in Africa and the Potential Emergence of Mosquito-Borne Arboviral Diseases—A Review and Research Outlook. In: Viruses.

Basnett, S., Krpan, J. & Espíndola, A. (2025). Floral traits and their connection with pollinators and climate. Annals of Botany, 135, 125–140.

Baude, M., Kunin, W.E., Boatman, N.D., Conyers, S., Davies, N., Gillespie, M.A.K. et al. (2016). Historical nectar assessment reveals the fall and rise of floral resources in Britain. Nature, 530, 85–88.

Blomberg, S.P., Garland, T.J.R. & Ives, A.R. (2003). Testing for Phylogenetic Signal in Comparative Data: Behavioral Traits Are More Labile. Evolution, 57, 717–745.

Bronstein, J.L., Alarcón, R. & Geber, M. (2006). The evolution of plant–insect mutualisms. New Phytologist, 172, 412–428.

Castellanos, M.C., Wilson, P. & Thomson, J.D. (2004). ‘Anti-bee’ and ‘pro-bird’ changes during the evolution of hummingbird pollination in Penstemon flowers. Journal of Evolutionary Biology, 17, 876–885.

Castro Tavares, D., Freitas, L. & Gaglianone, M.C. (2016). Nectar volume is positively correlated with flower size in hummingbird-visited flowers in the Brazilian Atlantic Forest. Journal of Tropical Ecology, -1, 1–5.

Corbet, S. (2003). Nectar sugar content: Estimating standing crop and secretion rate in the field. Apidologie, 34, 1–10.

Duchenne, F., Domínguez-García, V., Molina, F.P. & Bartomeus, I. (2026). Coevolution of phenological traits shapes plant-pollinator coexistence. Proceedings of the Royal Society B: Biological Sciences, 293, 20252808.

Farkas, Á. & Orosz-Kovács, Z. (2003). Nectar secretion dynamics of Hungarian local pear cultivars. Plant Systematics and Evolution, 238, 57–67.

Foster, W.A. (1995). Mosquito Sugar Feeding and Reproductive Energetics. Annual Review of Entomology, 40, 443–474.

Galetto, L. & Bernardello, G. (2004). Floral Nectaries, Nectar Production Dynamics and Chemical Composition in Six Ipomoea Species (Convolvulaceae) in Relation to Pollinators. Annals of Botany, 94, 269–280.

Gary Jr, R.E. & Foster, W.A. (2004). Anopheles gambiae feeding and survival on honeydew and extra-floral nectar of peridomestic plants. Medical and Veterinary Entomology, 18, 102–107.

Gouagna, L.C., Kerampran, R., Lebon, C., Brengues, C., Toty, C., Wilkinson, D.A. et al. (2014). Sugar-source preference, sugar intake and relative nutritional benefits in Anopheles arabiensis males. Acta Tropica, 132, S70–S79.

Gu, W., Müller, G., Schlein, Y., Novak, R.J. & Beier, J.C. (2011). Natural Plant Sugar Sources of Anopheles Mosquitoes Strongly Impact Malaria Transmission Potential. PLOS ONE, 6, e15996.

Harder, L.D. (1983). Effects of nectar concentration and flower depth on flower handling efficiency of bumble bees. Oecologia, 69.

Hicks, D.M., Ouvrard, P., Baldock, K.C.R., Baude, M., Goddard, M.A., Kunin, W.E. et al. (2016). Food for Pollinators: Quantifying the Nectar and Pollen Resources of Urban Flower Meadows. PLOS ONE, 11, e0158117.

Impoinvil, D.E., Kongere, J.O., Foster, W.A., Njiru, B.N., Killeen, G.F., Githure, J.I. et al. (2004). Feeding and survival of the malaria vector Anopheles gambiae on plants growing in Kenya. Medical and Veterinary Entomology, 18, 108–115.

Irwin, R.E., Bronstein, J.L., Manson, J.S. & Richardson, L. (2010). Nectar Robbing: Ecological and Evolutionary Perspectives. Annual Review of Ecology, Evolution, and Systematics, 41, 271–292.

Jin, Y. & Qian, H. (2019). V.PhyloMaker: an R package that can generate very large phylogenies for vascular plants. Ecography, 42, 1353–1359.

Kaczorowski, R.L., Gardener Mc Fau - Holtsford, T.P. & Holtsford, T.P. (2005). Nectar traits in Nicotiana section Alatae (Solanaceae) in relation to floral traits, pollinators, and mating system.

Keasar, T. & Bodner, L. (2025). Do flowers with specialized morphologies produce more nectar and pollen? American Journal of Botany, 112.

Kevan, P. & Baker, H. (1983). Insects as Flower Visitors and Pollinators. Annual Review of Entomology.

Lanuza, J.B., Rader, R., Stavert, J., Kendall, L.K., Saunders, M.E. & Bartomeus, I. (2023). Covariation among reproductive traits in flowering plants shapes their interactions with pollinators. Functional Ecology, 37, 2072–2084.

Lázaro, A., Vignolo, C. & Santamaría, L. (2015). Long corollas as nectar barriers in Lonicera implexa: interactions between corolla tube length and nectar volume. Evolutionary Ecology, 29, 419–435.

Manda, H., Gouagna, L.C., Foster, W.A., Jackson, R.R., Beier, J.C., Githure, J.I. et al. (2007). Effect of discriminative plant-sugar feeding on the survival and fecundity of Anopheles gambiae. Malaria Journal, 6, 113.

Martins, D.J. & Johnson, S.D. (2013). Interactions between hawkmoths and flowering plants in East Africa: polyphagy and evolutionary specialization in an ecological context. Biological Journal of the Linnean Society, 110, 199–213.

Müller, G.C., Junnila, A., Traore, M.M., Traore, S.F., Doumbia, S., Sissoko, F. et al. (2017). The invasive shrub Prosopis juliflora enhances the malaria parasite transmission capacity of Anopheles mosquitoes: a habitat manipulation experiment. Malaria Journal, 16, 237.

Müller, G.C., Xue, R.-D. & Schlein, Y. (2011). Differential attraction of Aedes albopictus in the field to flowers, fruits and honeydew. Acta Tropica, 118, 45–49.

Nyasembe, V.O., Cheseto, X., Kaplan, F., Foster, W.A., Teal, P.E.A., Tumlinson, J.H. et al. (2015). The Invasive American Weed Parthenium hysterophorus Can Negatively Impact Malaria Control in Africa. PLOS ONE, 10, e0137836.

Ornelas, J.F., Ordano, M., De-Nova, A.J., Quintero, M.E. & Garland Jr, T. (2007). Phylogenetic analysis of interspecific variation in nectar of hummingbird-visited plants. Journal of Evolutionary Biology, 20, 1904–1917.

Ortiz, P.L., Fernández-Díaz, P., Pareja, D., Escudero, M. & Arista, M. (2021). Do visual traits honestly signal floral rewards at community level? Functional Ecology, 35, 369–383.

Pagel, M. (1999). Inferring the historical patterns of biological evolution. Nature, 401, 877– 884.

Pattrick, J.G., Scott, J. & Wright, G.A. (2025). The concentration and energetic content of floral nectar sugars: calculation, conversions, and common confusions. Journal of Pollination Ecology, 38, 161–170.

Petanidou, T., Goethals, V. & Smets, E. (2000). Nectary structure of Labiatae in relation to their nectar secretion and characteristics in a Mediterranean shrub communityDoes flowering time matter? Plant Systematics and Evolution, 225, 103–118.

Petanidou, T. & Smets, E. (1995). The potential of marginal lands for bees and apiculture: nectar secretion in Mediterranean shrublands. Apidologie, 26, 39–52.

Revell, L.J. (2012). phytools: an R package for phylogenetic comparative biology (and other things). Methods in Ecology and Evolution, 3, 217–223.

Romero-Bravo, A. & Castellanos, M.C. (2024). Nectar and floral morphology differ in evolutionary potential in novel pollination environments. New Phytologist, 243, 753– 764.

Simpson, M.G. (2019). 9 - Plant Morphology. In: Plant Systematics (Third Edition) (ed. Simpson, MG). Academic Press, pp. 469–535.

Sinka, M., Pironon, S., Massey, N.C., Longbottom, J., Hemingway, J., Moyes, C. et al. (2020). A new malaria vector in Africa: Predicting the expansion range of Anopheles stephensi and identifying the urban populations at risk.

Smith, S.A. & Brown, J.W. (2018). Constructing a broadly inclusive seed plant phylogeny. American Journal of Botany, 105, 302–314.

Stone, C.M., Jackson, B.T. & Foster, W.A. (2012). Effects of plant-community composition on the vectorial capacity and fitness of the malaria mosquito Anopheles gambiae. Am J Trop Med Hyg, 87, 727–736.

Takkis, K., Tscheulin, T. & Petanidou, T. (2018). Differential Effects of Climate Warming on the Nectar Secretion of Early- and Late-Flowering Mediterranean Plants. Front Plant Sci, Volume 9–2018.

Temeles, E.J. & Dalsgaard, B. (2026). Specialization, generalization, and pollination syndromes: the role of pollinator competition. Evolutionary Ecology, 40, 2.

Tew, N.E., Memmott, J., Vaughan, I.P., Bird, S., Stone, G.N., Potts, S.G. et al. (2021). Quantifying nectar production by flowering plants in urban and rural landscapes. Journal of Ecology, 109, 1747–1757.

Torres, C. & Galetto, L. (2002). Are Nectar Sugar Composition and Corolla Tube Length Related to the Diversity of Insects that Visit Asteraceae Flowers? Plant Biology, 4, 360–366.

van der Kooi, C.J., Vallejo-Marín, M. & Leonhardt, S.D. (2021). Mutualisms and (A)symmetry in Plant–Pollinator Interactions. Current Biology, 31, R91–R99.

Wang, Y.H., Hsu, H.C., Chou, W.C., Liang, C.H. & Kuo, Y.F. (2020). Automatic Identification of First-Order Veins and Corolla Contours in Three-Dimensional Floral Images.

Xu, Q.-S. & Liang, Y.-Z. (2001). Monte Carlo cross validation. Chemometrics and Intelligent Laboratory Systems, 56, 1–11.

